# Discrimination of Gram-Positive versus Gram-Negative Bacteria by the Social Amoeba *Dictyostelium discoideum*

**DOI:** 10.1101/199471

**Authors:** Ghazal Rashidi, Elizabeth A. Ostrowski

## Abstract

Professional phagocytes detect, pursue, engulf, and kill bacteria. While all professional phagocytes use chemotaxis to locate bacteria, little is known about whether they can discriminate among them, responding preferentially to some bacteria over others. Here we examine the chemotaxis of the soil amoeba and professional phagocyte *Dictyostelium discoideum* in assays where amoebae were presented with a paired choice of different bacteria. We observed variation in the extent to which they pursue different types of bacteria and preferential migration towards Gram-negative over Gram-positive bacteria. Response profiles were similar for amoebae isolated from different geographic locations, suggesting that chemotaxis preferences are not strongly influenced by any local variation in the bacterial community. Because cyclic adenosine monophosphate (cAMP) is a known chemoattractant for *D. discoideum*, we tested whether it mediated the preference for Gram-negative bacteria. Chemotaxis was diminished in response to a cAMP-deficient strain of *Escherichia coli* and enhanced in response to an *E. coli* strain that overproduces cAMP. We conclude that *D. discoideum* discriminates at a distance among bacteria and that discrimination is mediated in part by sensing of cAMP. Preferential sensing and response to different bacteria may help to explain why some amoeba-bacterial associations are more prevalent in nature than others.

## Background

There is a growing appreciation that nearly all multicellular organisms are colonized by large numbers of bacteria that approach or even outnumber an organism’s own cells [1–3]. These bacteria can influence everything from a host’s physiology to its development and behavior [reviewed in 4–6]. Despite the near universality of colonization by bacteria, however, little is known about the extent to which hosts can selectively maintain or remove different bacteria— which in turn requires the ability to sense and respond preferentially to some bacteria over others.

Professional phagocytes include motile cells of the mammalian immune system, such as neutrophils or macrophages, that locate and engulf bacterial invaders [7–9]. However, phagocytosis is also used by free-living amoebae to hunt and kill bacteria for food [10,11]. Phagocytosis involves invagination of the cell membrane, engulfment, and internalization of a bacterial cell or object. The resulting vacuole, called a phagosome, subsequently fuses with a lysosome that contains a variety of hydrolytic enzymes and usually results in death and digestion of the bacteria. Because the molecular basis of phagocytosis is conserved between amoebae and mammals, many bacteria that can evade intracellular killing and replicate inside immune phagocytes also infect and kill free-living amoebae [12,13]. Indeed, amoebae are thought to be the primary reservoir for these bacterial pathogens in nature and potentially a natural predator that selected for their pathogenic features [14–20].

Phagocytosis is a specific process, requiring host receptors that recognize and bind ligands on the bacterial cell surface [21–25]. This specificity alone leads to some degree of discrimination on the part of phagocytes in their ability to attach to, engulf, and kill bacteria they encounter [26]. In the soil amoeba *Dictyostelium discoideum*, for example, distinct sets of genes are required for killing and digestion of Gram-negative versus Gram-positive bacteria, and different genes respond transcriptionally to phagocytosis of these two classes of bacteria [27].

In addition to binding and engulfing bacteria once they have been encountered, professional phagocytes show a variety of chemotactic or “chase” behaviors [28] that enable them to detect and respond to bacteria at a distance (e.g., neutrophil chase [29]). In some cases, the bacteria- or host-produced chemoattractants have been identified. For example, N-formylated peptides, produced by bacteria, are chemoattractive to neutrophils [30,31]. Macrophage and neutrophils are also attracted to components of the complement system that are released by other cells upon contact with bacteria [32–34].

Despite work on immune phagocyte discrimination upon physical contact, there have been few attempts to demonstrate that phagocytes can discriminate among different types of bacteria at a distance, allowing them to selectively pursue certain bacteria, despite the importance of any such discrimination for outcomes such as clearance, infection, or colonization. In one notable exception, previous work showed that neutrophils will extend more pseudopods towards *E. coli* and *Salmonella enterica* serovar *Typhimurium* than they do towards *S. enterica* serovar *Typhi*. It was also shown that expression of the Vi capsular polysaccharide by the latter impedes neutrophil chemotaxis and leads to the discriminatory response [35].

*Dicytostelium discoideum* is soil amoeba and major model system for phagocytosis, chemotaxis, and host-pathogen interactions. As a single-celled amoeba, it phagocytoses bacteria for food during the unicellular stage of its life cycle. In response to starvation, it undergoes aggregative multicellular development, mediated by cyclic AMP signaling, first forming a migratory slug, and later a fruiting body that consists of a ball of spores atop a stalk of dead cells. Many bacteria that infect and kill macrophage or neutrophils also infect and kill *Dictyostelium* [12,36–39], suggesting that bacteria can function as both predator and prey to *D. discoideum*. Early studies by Konijn and colleagues [22] demonstrated that *D. discoideum* will respond chemotactically to *E. coli* and possibly other species of bacteria, but whether the amoebae discriminate among different bacteria and respond selectively is not known. Moreover, while it was long thought that the multicellular stages of the life cycle (the slug and fruiting body) were sterile, a recent study showed that some *D. discoideum* strains are colonized by bacteria [40]. Additional work demonstrated variable impacts of these colonizing bacteria on the amoebae that range from beneficial to commensal [41–43]. These diverse impacts of bacteria on amoebae, in addition to its reliance on bacteria for food, makes *D. discoideum* a useful model system to test for the possibility of discrimination at a distance.

To determine whether *D. discoideum* can discriminate among different bacteria, we performed behavioral choice assays, where amoebae were placed equidistant between different species of bacteria, and we quantified the movement by the amoebae to each one. Our experiments consisted of pairwise combinations of four Gram-negative and four Gram-positive species. We chose Gram-negatives *Klebsiella pneuomoniae* and *E. coli* because these bacteria are high quality prey for *D. discoideum*, which can devour them rapidly and develops well on lawns of both species. Gram-negatives *Pseudomonas fluorescens* and *Burkholderia* spp. have both been repeatedly isolated from *D. discoideum* fruiting bodies [40], and we hypothesized that the amoebae might show greater sensitivity to, or even preference for, these bacterial species. Indeed, the particular isolate of *Burkholderia* we used was isolated from a fruiting body of a *D. discoideum* natural isolate in the lab. Among the Gram-positives, *Bacillus subtilis* is a common soil bacterium that *D. discoideum* might encounter. *Microccocus luteus* is likewise common in the environment, but also found on skin. *Enterococcus facaelis* is a Gram-positive enteric bacteria that *Dictyostelium* could potentially encounter in nature, since naturally occurring *D. discoideum* fruiting bodies have only been found on dung [e.g., 45]. Finally, *Staphylococcus aureus* is a Gram-positive bacterium commonly found on the skin and in the respiratory tract. Overall, we chose a range of bacteria that likely vary with respect to the extent of *D. discoideum*’s prior experience with them during its evolutionary history.

Finally, we tested three different *D. discoideum* genotypes from distinct geographic regions (Massachusetts, Texas, and Tennessee) that comprise a large portion of the species’ natural range, which is primarily the eastern half of the United States, as well as parts of central America and east Asia [45]. The degree to which chemotaxis behaviors are similar or different among geographically and genetically diverged *D. discoideum* can help to determine whether bacteria-seeking behaviors are evolutionarily malleable and responsive to natural variation in the bacterial community across environments.

## Methods

### Dictyostelium Strains and Culture Conditions

*D. discoideum* strains QS32 (Houston, Texas), QS39 (Indian Gap, Tennessee), and QS40 (Mt. Mitchell, Massachusetts) were spotted from frozen spore stocks onto lawns of *Klebsiella pneumoniae* bacteria on SM agar plates (per liter: 10 g glucose, 10 g Bacto Peptone (Oxoid), 1 g yeast extract (Oxoid), 1g MgSO_4_, 1.9 g KH_2_PO_4_, 0.6 g K_2_HPO_4_, 20 g agar). Following germination and growth, amoebae were harvested from a leading edge and inoculated into petri dishes containing 10 ml of HL5 medium (ForMedium Ltd) supplemented with 10% fetal bovine serum and 2% PSV (10 mg/ml penicillin, 50 mg/ml streptomycin sulfate, 20 μg/ml folate, and 60 μg/ml vitamin B12). Cells were maintained as axenic (bacteria free) adherent cultures for approximately one week prior to the start of the experiment.

### Bacterial Strains and Culture Conditions

Eight bacterial species were streaked onto lysogeny broth (LB) agar plates from frozen cultures: Gram-negative: *K. pneumonaie (Kp)*, *E. coli (Ec)*, *P. fluorescens (Pf)*, *Burkholderia spp. (Bu)*; Gram-positive: *E. facaelis* (Ef), *B. subtillus* (*Bs*), *S. aureus* (*Sa*), or *M.*(*Ml*). Strain information is provided in Table S1; most were standard lab strains, except for *Burkholderia* spp., which we isolated from the sorus of a *D. discoideum* fruiting body. Following inoculation from frozen stocks, colonies were inoculated into 1 ml of LB medium and grown overnight at their respective optimal temperatures: 37C (*Ec*, *Bs*, *Kp*, *Ml*, and *Sa*), 31C (*Bu*, *Ef*) or room temperature (*Pf*).

### Chemotaxis Assay

Following overnight growth, bacterial cultures were concentrated to an OD_600_ of 5 by centrifugation. Cells of *D. discoideum* were harvested from the petri dishes by pipetting, washed with KK2 buffer (14.0 mM K_2_HPO_4_ and 3.4 mM K_2_HPO_4_, pH = 6.4), and re-suspended at a density of 1 × 10^7^ cells/ml. Two microliters of each *D. discoideum* suspension (corresponding to 2 × 10^4^ cells) were spotted onto a Nunc Omni tray containing 10 ml of KK2 with 2% agar noble. A grid was placed beneath the plate to ensure equal spacing of the amoeba spots, which were placed at distances of approximately 1.3 cm. The *D. discoideum* spots were first allowed to dry (~15 min), and then 2 μl spots of bacteria were placed in between each *D. discoideum* spot, again using a grid beneath the plate for placement, such that the distance between the center of each *D. discoideum* spot and those of its bacterial choices was approximately 0.65 cm (see Figure 1). Each *D. discoideum* genotype was placed in a different row of the plate, and each bacterial species in a different column. Across replicate experiments, we changed the order of column and rows so that the bacteria and *D. discoideum* cells were in different locations. Experiments were carried out in three temporally independent blocks, each commenced by re-inoculating from frozen stocks.

**Figure 1.**
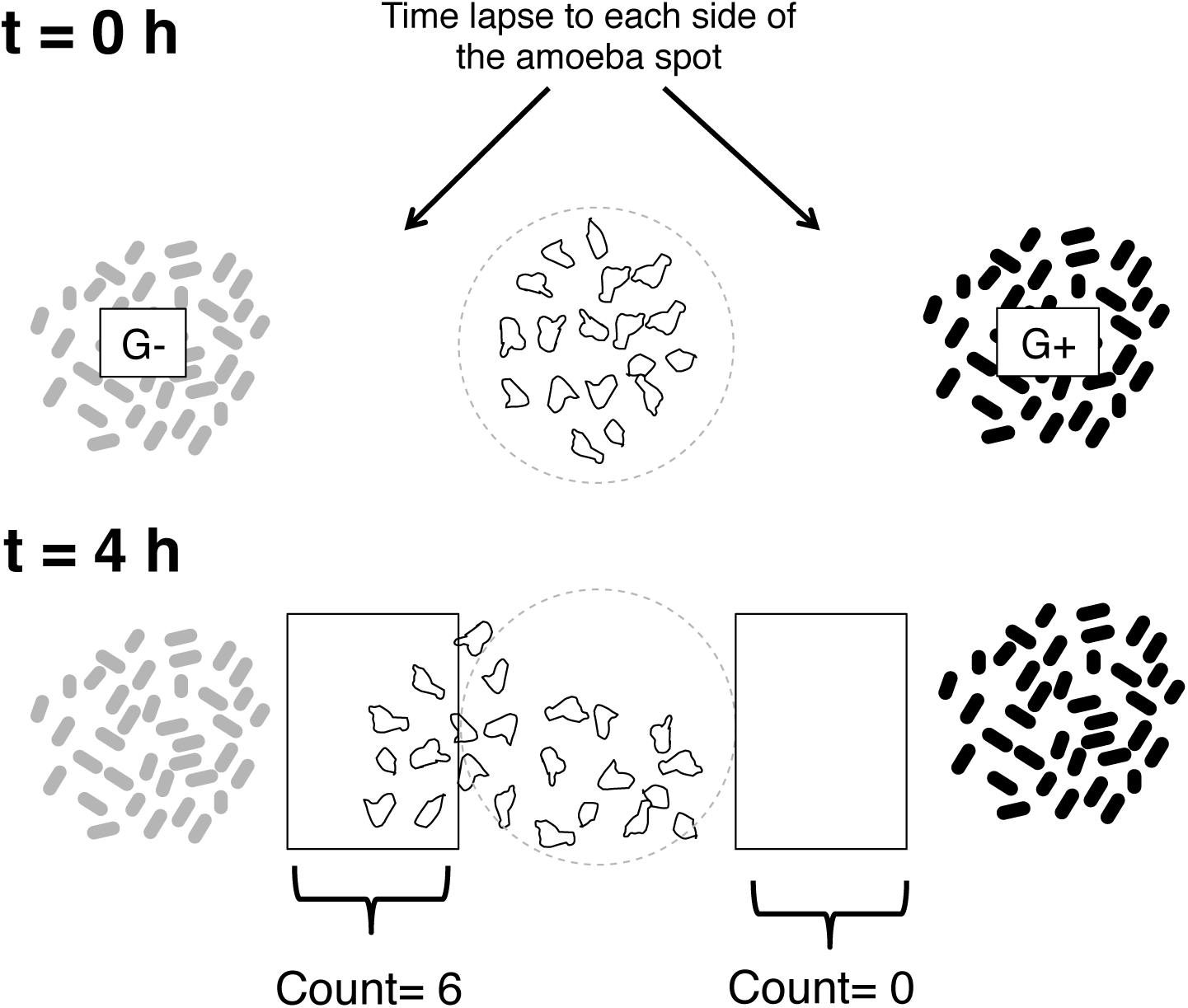
Experimental design for paired choice assay. Amoebae of each of three *D. discoideum* genotypes (QS32, QS39, or QS40) were deposited in spots on a plate containing a thin layer of agar, equidistant between spots of different pairs of Gram-negative and Gram-positive bacteria. A given *D. discoideum* strain was spotted across a row, and a given bacterial species was spotted down a column, creating different combinations of *D. discoideum* genotypes and paired choices of Gram-negative and Gram-positive bacteria. The order of the *D. discoideum* genotypes in rows, and the order of the Gram-negative and Gram-positive bacteria in columns, was varied across replicate experiments. The left and right edge of each amoeba spot was recorded by time-lapse video, and the image corresponding to 4 hours was extracted for analysis. The 4 hour timepoint represented a point where the amoebae had consistently begun migrating toward the bacteria but typically had not yet filled the entire field of view. We quantified the number of amoebae present in the image after cropping it at a tangent to the original spot.

### Time Lapse Videos of Chemotaxis

We recorded the movement of the *D. discoideum* cells toward the bacteria by time-lapse microscopy, which commenced two hours after the spots were deposited. Images were taken on a Leica DMI6000B microscope with an automated stage at 50X magnification using the LAS software suite. The left and right edges of each amoeba spot (representing the area the amoebae would enter once they commenced chemotaxis) were imaged every five minutes. From these image stacks, we extracted the image corresponding to four hours from the start of the experiment, which was two hours after the start of the time lapse. We chose this time-point based on preliminary experiments: by this time, the amoebae had usually exited the boundaries of the spots, but not yet filled the imaged area completely.

### Image Analysis

The number of cells that moved towards each bacterial species was counted in ImageJ. Images were first cropped (at a tangent to the spot) so that only cells that had migrated outside the boundaries of the spot would be counted. The cropped image, consisting of a rectangular selection of dimensions 10 × 12.5, was converted to an 8-bit image, thresholded to achieve a binary image, and the cells were counted using the module “Analyze Particles”. Some manual adjustment was necessary to ensure clumps were split into single cells and any holes in the cells were filled. See Fig. S1 for an example of a raw image and the same image following thresholding and counting.

### Adenlyate Cyclase E. coli Mutants

To assess the impact of cAMP on *D. discoideum* chemotaxis, we obtained a cAMP-deficient *E. coli* strain, where the only adenylate cyclase gene (*cyaA*), which is necessary for cAMP production, had been deleted (Keio:JW3778; *∆cyaA*::kan^R^ allele) and its wild-type progenitor (BW25113) from GE Dharmacon [46]. We also obtained an *E. coli* strain that carries an additional copy of the *cyaA* gene on an inducible plasmid (*cyaA^OE^;* ASKA *E. coli* K12 W3110; gift of Tim Cooper, University of Houston) [47]. The presence and absence of the *cyaA* gene in the *∆cyaA* and wild-type *E. coli*, respectively, was confirmed by PCR using *cyaA* specific primers, whereas the ASKA strain was confirmed by sequencing the plasmid insert.

Prior to the assays, the *∆cyaA*, wild-type, and *cya^OE^* strains were streaked onto LB plates, grown overnight at 37 C, and single colonies were inoculated into test tubes containing 1 ml LB. The *∆cyaA* culture was supplemented with 30 μg/ml kanamycin to ensure retention of the deletion allele, and the *cyaA*^OE^ culture was supplemented with 10 μg/ml chloramphenicol and 1 mM IPTG to promote plasmid maintenance and induce expression of the *cyaA* gene, respectively. All cultures were incubated overnight at 37 C and concentrated to an OD_600_ of 5 prior to the start of the experiment. Chemotaxis assays were carried out as described above. We used all possible pairwise combinations of the three bacterial genotypes, with replicate spots on the plate for each *D. discoideum* - *E. coli* genotype pair. The entire assay was then repeated in a temporally independent replicate.

### Statistical analysis

The experiment results were analyzed as a mixed-model ANOVA using PROC MIXED in SAS (v. 9). The number of cells migrating after 4 hours was modeled as a function of the *D. discoideum* (Dd) genotype (QS32, QS39, or QS40), the focal bacterial species, the alternative bacterial choice, and the block (i.e., experimental replicate). Focal bacteria and alternative bacteria were nested within Gram-status (positive or negative). We included interaction terms in the model, Dd genotype × bacteria, Dd genotype × alternative bacteria, and Dd genotype × block. The significance of random effects factors was assessed using a likelihood ratio test that compared the difference in the -2 residual likelihoods of nested models that did or did not include the factor of interest. This difference was compared to the Chi-square distribution, with the degrees of freedom equal to the difference in the number of parameters between the full and reduced models. Gram status was modeled as a fixed effects factor, and the significance of this term was determined by a standard F-test.

## Results

In 21 of 24 pairwise choices between a Gram-positive and a Gram-negative bacterium, *D. discoideum* preferentially migrated towards the Gram-negative bacterial species (Fig. 2; sign-test: two-tailed *P*<0.0003). There were three exceptions to this pattern: QS39 showed a weak preference for Gram-positive *B. subtilis* over Gram-negatives *K. pneumonaie* and *E. coli*, and QS40 weakly favored *B. subtilis* in a pairwise contest against *K. pneumonaie* (see Fig. 2).

**Figure 2.**
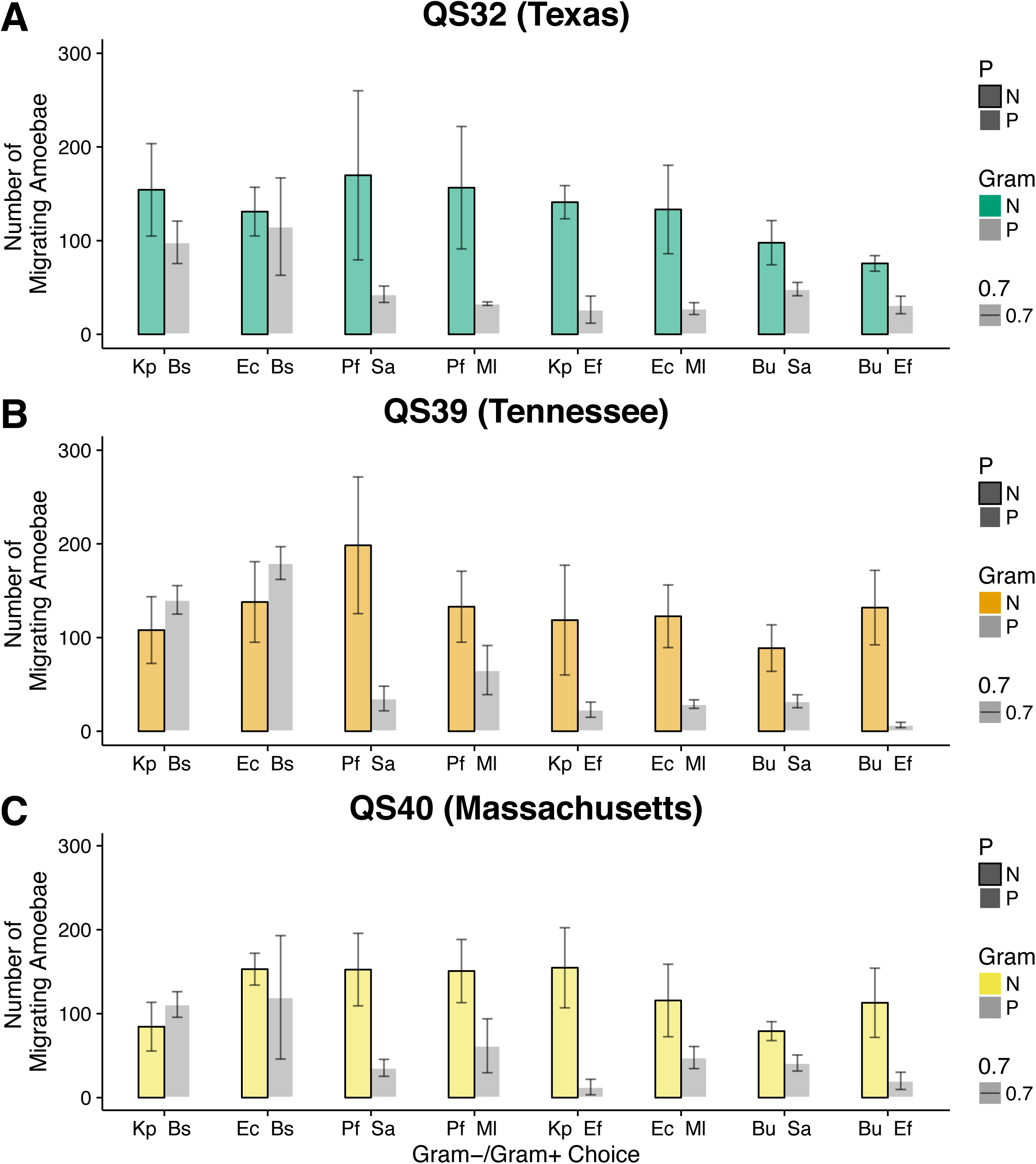
Pairwise choice assays reveal preference for Gram-negative over Gram-positive bacteria. Each paired bar shows a choice assay between the two indicated spcies of bacteria, with the Gram-negative bacteria plotted on the left, and the Gram-positive bacteria on the right. Error bars represent one standard error across temporally independent replicates of the experiment. In 22 of 24 assays, more cells migrated towards the Gram-negative compared to the Gram-positive bacteria.

We modeled the chemotactic response, quantified as the number of cells migrating towards a bacterial spot after 4 hours, as a function of the focal bacteria, the alternative choice of bacteria (see Fig. 1), the Gram status, and the *D. discoideum* genotype. All three interaction terms (Dd genotype × focal bacteria, Dd genotype × alternative bacteria, and alternative bacteria × focal bacteria) were highly non-significant and dropped from the model (Table 1). The identity of the alternative bacteria in each pairwise choice was also highly non-significant and likewise dropped from the model. The lack of significance for this term indicates that the chemotaxis of the amoebae to any given bacteria was similar regardless of its alternative choice bacteria.

**Table 1.** Results of mixed model analysis of variance for effects of random factors. Each random factor was tested by taking the difference between the -2 residual log likelihoods of nested models with and without the factors of interest, with degrees of freedom equal to the difference in the number of parameters between the nested models. The only fixed effect factor, Gram status, was tested using a standard F-test (*F*_1,5.87_=9.23, *P*=0.02), as recommended (Bolker et al. 2009)

| Model | -2 Residual Log Likelihood (-2RLL) | Difference in -2RLL ($\sim\chi^2$) | Difference in model d.f. | *P* |
| --- | --- | --- | --- | --- |
| Full Model | 1822.9 |  |  |  |
| Reduced Models, Removing: |  |  |  |  |
| Interaction Terms | 1823.4 | 0.5 | 3 | 0.97 |
| *Dd* Genotype | 1823.4 | 0* | 1 | 1 |
| Alternative Bacteria | 1823.4 | 0* | 1 | 1 |
| Block | 1846.0 | 22.6 | 1 | <0.0001 |
| Focal Bacteria | 1852.4 | 29 | 1 | <0.0001 |

\*No change in -2 residual log-likelihood when removing this factor. We obtained a similar, non-significant effect of *Dd* genotype or alternative bacteria when using a least-squares approach.

Overall, we observed significant variation in the response to different bacteria (focal bacteria; χ^2^=29, df=1, *P*<0.0001). The strongest response was to *P. fluorescens*, a bacterial species that has been found in the fruiting bodies of *D. discoideum* [40,42]. The weakest chemotactic responses were to the Gram-positive bacteria *M. luteus*, *S. aureus*, and *E. facaelis* (Fig. 3).

**Figure 3.**
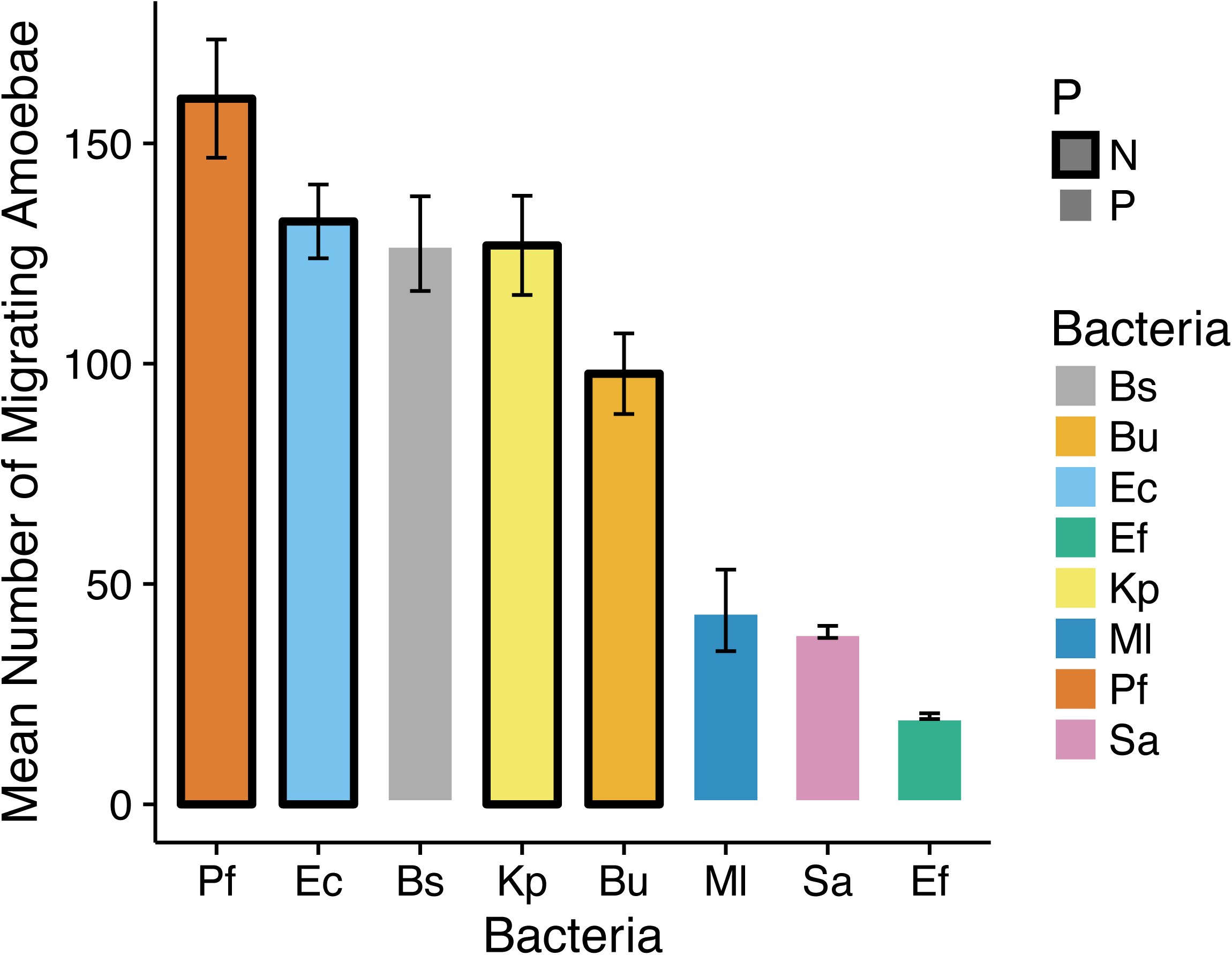
Mean response to each bacterial species, ordered from strongest response to weakest response. Each bar shows the grand mean response to each bacterial species, averaged across the three *D. discoideum* genotypes. Dark borders indicate Gram-negative bacteria, while light borders indicate Gram-positive bacteria.

Consistent with our conclusions based on the pairwise analysis, our full statistical model showed a significant effect of Gram-status on chemotaxis (Fig. 4; *F*_1,5.87_=9.23, *P*=0.02), which reflected an overall preference for Gram-negative bacteria. Despite some differences among *D. discoideum* genotypes in the level of response to *B. subtilis*, the overall response profile to different bacteria was strikingly similar across the geographically widespread *D. discoideum* strains (Table 1, non-significant effect of *Dd* genotype of chemotaxis: χ^2^ =0, df=1, *P*=1).

**Figure 4.**
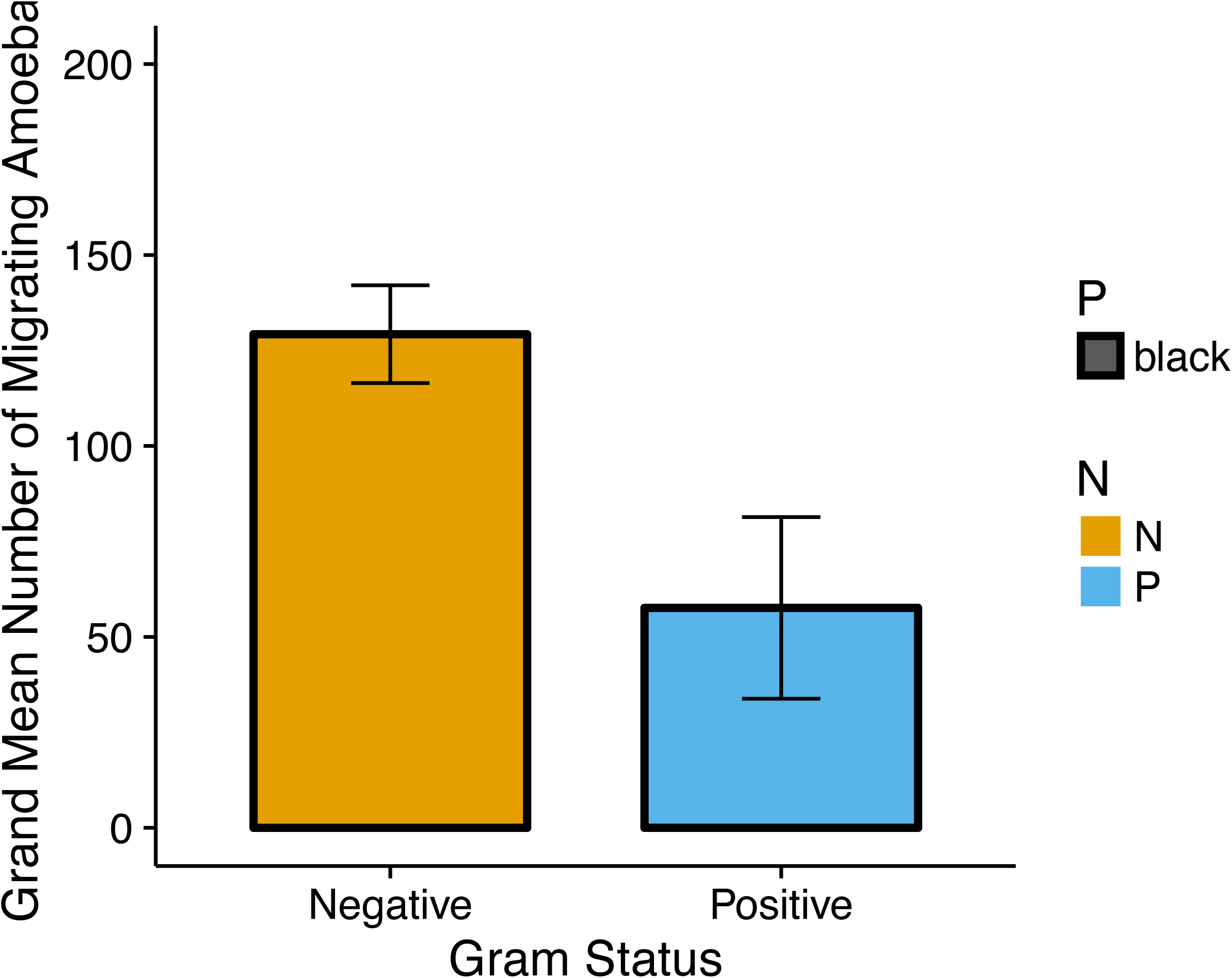
Mean response to Gram-negative versus Gram-positive bacteria. ANOVA results indicate that amoebae showed significant preference for Gram-negative bacteria (P=0.02; see text for details.)

Why does *D. discoideum* exhibit a chemotactic preference for Gram-negative bacteria? We hypothesized that this effect might be mediated by cAMP, a known chemoattractant for *D. discoideum* that is also differentially produced by Gram-negative and Gram-positive bacteria. In Gram-negative bacteria, cAMP functions in catabolite repression and is produced in large quantities intracellularly and released into the medium when glucose is absent. In contrast to the central role of cAMP in Gram-negative bacteria, Gram-positive bacteria do not use cAMP to mediate catabolite repression, there are few reports of its detection in Gram-positive bacteria except under limited conditions (e.g., O2 limitation [48]), and there are no established signaling roles for cyclic mononucleotides in Gram-positive bacteria [49].

To test the hypothesis that amoebae are specifically sensing and responding to cAMP, we compared the amoeba response to an *E. coli* mutant with a gene deletion of *cyaA*, which encodes adenylate cyclase, the enzyme that catalyzes the production of cAMP from adenosine triphosphate (ATP). We compared the amoebae responsiveness to the deletion mutant to that of its wild-type progenitor and to an *E. coli* strain that overexpresses *cyaA*—and thus these strains should vary in their production of cAMP. Consistent with our hypothesis, chemotaxis was strongest to *cyaA^OE^* and weakest to *ΔcyaA* (see Fig. 5 for example images for strain QS39), and the pattern was consistent across all three *D. discoideum* genotypes (Fig. 6). Taken together, this indicates that the positive response to *E. coli* is mediated, at least in part, by sensing of bacterially produced cAMP.

**Figure 5.**
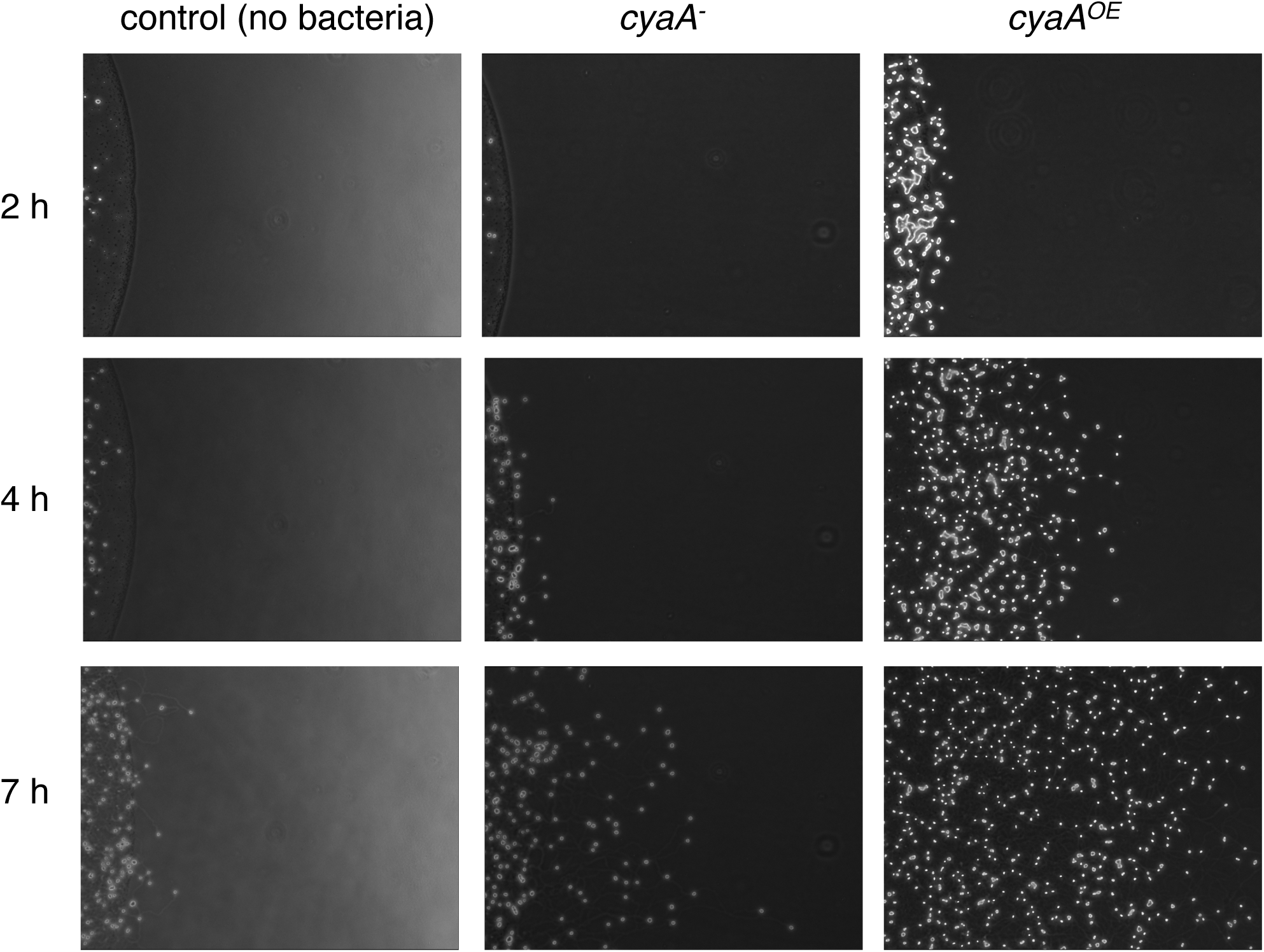
Example migration results for QS39. Columns show the time progression of amoeba migration out towards the indicated bacterial strain at 2 h, 4h, or 7 h following the start of the experiment.

**Figure 6.**
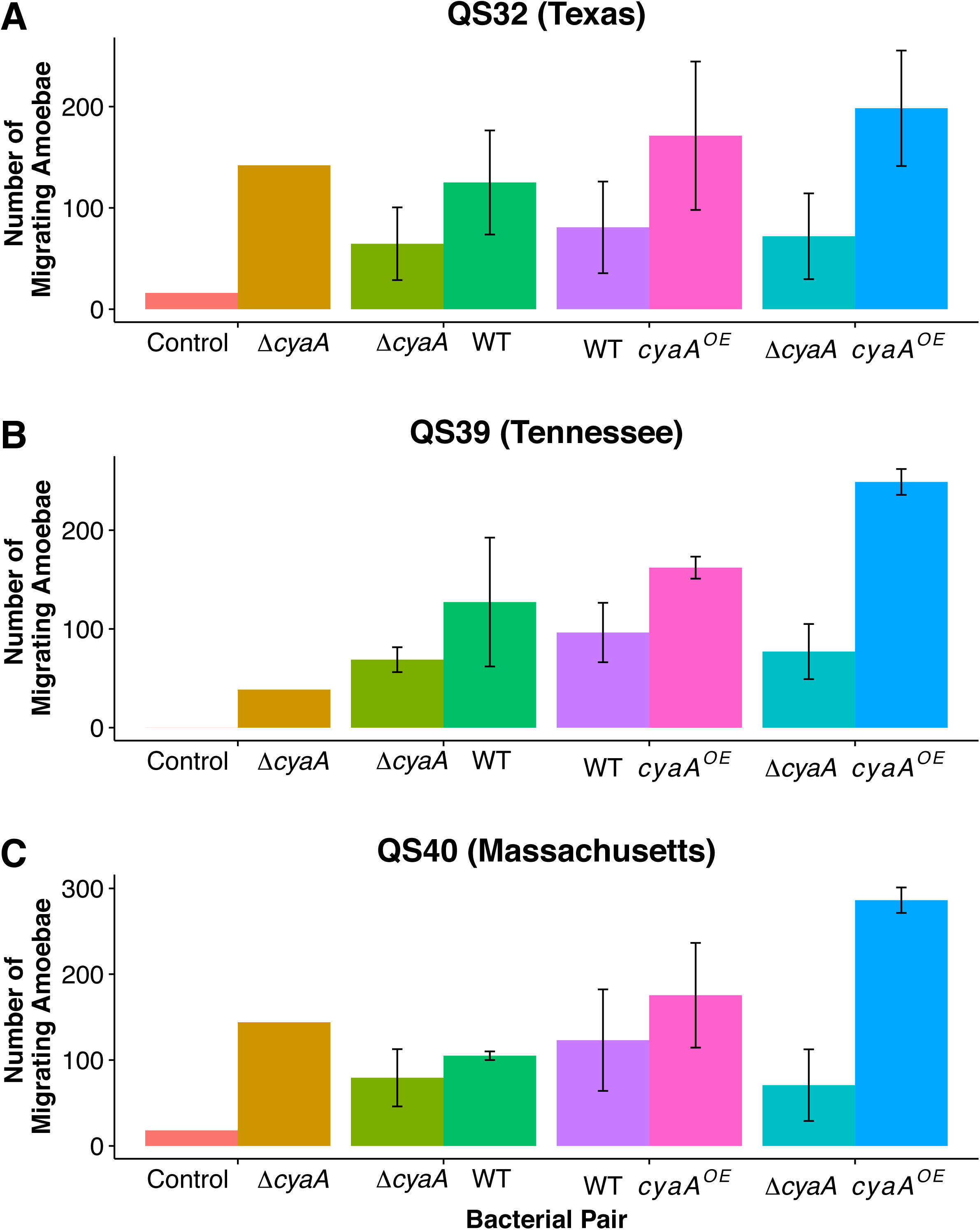
Migration towards *E. coli* strains that vary in cAMP production. We compared the response to a wild-type E. coli, and two mutants *ΔcyaA* and *cyaA*^OE^, where the adenylate cyclase gene necessary for production of cAMP is deleted or overexpressed, respectively. Consistent with our hypothesis, there was stronger migration in each paired comparison to the *E. coli* strain producing more cAMP. Note, however, that there was also stronger migration to the *ΔcyaA* strain than in controls containing no bacteria.

## Discussion

Professional phagocytes locate bacteria in part based on signals secreted by bacteria. The ability to find bacteria at a distance is likely crucial for amoebae that seek bacteria for food and phagocytic immune cells that must arrive at a site of infection and mount an immune response or else allow a beneficial symbiont to remain and colonize. In the social amoeba *D. discoideum*, environmental bacteria represent not only potential prey, but some bacteria can establish intracellular infections that range from pathogenic to mutualistic [11,40–42,50,51]. Discrimination may be critical to explain why some bacteria are readily localized and removed by phagocytes whereas others evade detection.

In paired choice assays, we observed variation in the responsiveness of amoebae to different bacteria, and the overall pattern was stronger in favor of Gram-negative compared to Gram-positive bacteria. While it is known that *D. discoideum* can respond to folate derivatives released by bacteria [52,53] and it has been suggested previously that cAMP might mediate bacterial sensing [54,55], no prior work has quantified or demonstrated differential chemotaxis that effectively discriminates among different types of bacteria.

We initially hypothesized that, because chemotaxis towards bacteria is likely an adaptive response by amoebae to locate potential food sources, we might observe different chemotactic profiles of *D. discoideum* isolates from different geographic locations where the local bacterial community may vary. While we did observe some differences in response to the Gram-positive *B. subtilis*, overall we found no significant difference among the three geographically divergent D. *discoideum* genotypes in their bacterial chemotaxis profiles. This result suggests that discrimination ability may relate to fundamental characteristics of the bacteria, in that it is not highly malleable to fine-scale differences in the composition of the bacterial community across sites.

In light of cAMP as a strong chemoattractant for *D. discoideum* during its developmental transition to multicellularity and also its differential use by Gram-negative and Gram-positive bacteria, we examined the potential role of this molecule in mediating the chemotaxis behavior of *D. discoideum*. Consistent with earlier work, all three amoeba genotypes responded as expected under our hypothesis: their chemotaxis response was strongest to the *cyaA*^OE^ (which should produce the highest levels of cAMP), weaker towards the wild-type, and the lowest to *cyaA* null mutants that lack cAMP. Taken together, this result suggests that *D. discoideum*’s preference for Gram-negative bacteria is mediated in part by differences in production of cAMP among bacteria. We note, however, that we observed stronger migration towards the *cyaA-* strain than a control that lacked any bacteria, indicating that cAMP is not the sole molecule that induces chemotaxis towards *E. coli*. The likely explanation for the migration towards the *cyaA-* strain is that the cells are responding to folate, which has been previously associated with localization of bacteria by *Dictyostelium* [52,56], or some other unidentified chemoattractant.

The specific bacteria that induced the strongest amoeba response are also interesting to consider individually. *P. fluorescens*, which elicited the strongest overall response, has been repeatedly isolated from the fruiting bodies of *D. discoideum* and shown to provide some benefit to the amoeba, as either food or through the production of antifungal compounds [42]. Two other bacteria that also elicited strong responses were *E. coli* and *K. pneumoniae* – these are both species that are routinely used to cultivate *D. discoideum* in the laboratory and constitute the best known prey bacteria for this species.

In prior work, *Burkholderia, Enterobacter sakazakii*, and *P. fluorescens* were commonly isolated from *D. discoideum* sori [40], and our characterization of the bacterial soil community at sites in Virginia where *D. discoideum* is unusually prevalent also indicates that these species are common in the soil (unpublished results). Taken together, these findings could suggest that *D. discoideum* may be most capable of detecting the bacteria with which it associates or encounters frequently in nature. However, the directionality of this finding is not obvious. In other words, does *D. discoideum* become preferentially associated with these bacteria because it readily detects and finds them in nature? Or has its detection method been fine-tuned to preferentially sense and find the bacteria with which it best associates? While we cannot decisively answer this question, the lack of any geographic pattern in the response of *D. discoideum* genotypes to different bacterial species, as well as the potential role of cAMP as the molecule that *D. discoideum* senses, lends greater credence to the first hypothesis – that *D. discoideum* is biased towards finding and associating with certain bacteria because of the conserved mechanism by which it senses them. More generally, our results suggest that long-term colonization, particularly of professional phagocytes, might be mediated in part by the mechanisms by which they detect and respond to bacteria and suggests the possibility that hosts might actively pursue and determine colonization outcomes.

## Author Contributions

GR carried out experiments and analyzed data. EAO carried out experiments, performed statistical analyses, and wrote the paper. All authors gave final approval for publication.

